# On Temporal Robustness & Brain-State Stability of Functional Connectivity in Mouse Primary Visual Area V1 compared to Higher Visual Area AL

**DOI:** 10.1101/2025.11.10.687557

**Authors:** Mario Alexios Savaglio, Christina Brozi, Stelios M. Smirnakis, Maria Papadopouli

## Abstract

Understanding how the structure of functional connectivity in the visual cortex changes over time and across brain states is crucial for elucidating the mechanisms by which neurons coordinate to process information and support behavior. Higher-order visual areas in mice are known to exhibit more distinct, segregated functional roles compared to the primary visual cortex (V1) [1], and they maintain stimulus representations over extended time scales [2]. However, the stability of the architecture of their functional connectivity across time and brain states remains less understood. In vivo mesoscopic two-photon calcium imaging was used to simultaneously record activity from thousands of neurons across V1 and the extra-striate anterolateral area (AL) in mice, during both visual stimulation (optical-flow/motion) and at resting state (i.e., absence of stimulus). We then applied the spike time-tiling (STTC) coefficient [3] to estimate the pairwise correlations of the neuronal firing and form the *functional connectivity* at the cell resolution. We then comparatively analyzed the functional connections within area AL and V1 under both stimulus-driven and resting-state conditions. The functional connectivity within AL remains consistently more robust over time than in area V1. Moreover, the structure of the functional connectivity in AL exhibits a smaller change between these two conditions compared to V1, indicating that functional connectivity derived from spontaneous activity more faithfully reflects the functional network architecture elicited by visual stimulation in this higher-order area. Finally, during the resting state, AL activity and functional connectivity are less dependent on pupil size than those of V1, indicating that arousal exerts a weaker modulatory effect on AL compared to V1.

## I. Introduction

In the visual cortex, signals traverse a hierarchy of processing stages—from primary visual cortex (V1) to higher visual areas (HVAs) [4]—each responsible for extracting distinct and increasingly abstract features [1], [5]–[7]. Harris *et al*. [8] assigned a hierarchy score to 37 cortical and 24 thalamic regions in mice based on their layer-specific axonal termination patterns. This revealed a hierarchical ordering of visual areas. The primary visual area V1 was found to be the lowest cortical area in the hierarchy, while other visual cortical regions, such as the anterolateral area (AL), were assigned higher hierarchy scores. Piasini *et al*. [2] observed that the neural responses in HVAs decay more slowly over time than those in V1.

Functional connectivity can reveal coordinated processing through temporally synchronized neuronal activity and has been discussed in numerous previous works, e.g., [9]–[14]. Siegle *et al*. [7] found that the organization of inter-areal functional connectivity during visual stimulation mirrors the anatomical hierarchy established by Harris *et al*. [8]. Additionally, Yu *et al*. [15] analyzed the functional connectivity both within and across cortical areas and found that, under visual presentation, pairwise noise correlations are more stable across different types of stimuli (moving gratings and naturalistic videos) compared to signal correlations. However, comparative analysis of the stability of the intra-areal functional connections between different cortical areas across time, and especially across distinct brain states, remains largely unexplored.

Population activity and functional connectivity during spontaneous conditions in mice V1 can be modulated by arousal [14], [16]. Additionally, neuronal responses in certain higher-order areas have been shown to be less modulated by arousal compared to those in V1 [17]. However, it is unclear how dependent the functional connectivity of higher visual circuits is on factors, such as the presence of visual input or varying arousal states, compared to that of V1. Addressing this gap will provide insights into how neural processing in V1 and HVAs is affected by the visual stimulus and brain state.

We used *vivo mesoscopic 2-photon calcium imaging* to record essentially simultaneously from *thousands* of pyramidal neurons in mouse area V1 and higher-order visual areas, under both visual stimulation and resting-state conditions. Here we focus on granular (L4) and supragranular (L2/3) layers of area V1 and the higher-order anterolateral area (AL). Images were appropriately thresholded to yield calcium “eventograms” per neuron, a proxy of the firing events (see Section II, Appendix). These binary time series were analyzed. The pairwise correlation metric Spike Time Tiling Coefficient (STTC) was applied on the eventograms [3] to form the functional connectivity of the circuits in V1 and AL under both visual stimulation and resting state, focusing on the *intralayer* connections within L2/3 and L4. To assess temporal robustness, we computed the number of functional connections that are persistent across time. We then compared the structure of the functional connectivity network derived under visual stimulation with that observed during the resting state. We also examined how the size of the pupil, a proxy for arousal [16], [19], modulates the activity and functional connectivity of each area, quantifying the stability of the functional connections to changes in brain state. Finally, we assessed the degree to which our observations are dependent on inter-neuronal distance. We found that compared to V1, the functional connectivity architecture of AL exhibits higher temporal robustness and brain-state stability. The rest of this paper is structured as follows: Section II overviews the experiments, the datasets, and the neural population. In Section III, we characterize the functional connectivity in both V1 and AL. Section IV assesses the stability of the functional connections within and across the visual presentation and resting-state conditions. In Section V, we present how arousal modulates the activity and functional connectivity during the resting state, while Section VI examines the potential spatial locality bias. Finally, Section VII discusses our main results and future work plans.

## II. Experiments, Data Collection, and

### Pre-Processing

This study focuses on data collected from the granular (L4) and supragranular (L2/3) layers (Fig. 1A) within the primary visual cortex (V1) and the anterolateral (AL) higher-order visual area (Fig. 1B) in five adult mice.

**Fig. 1:**
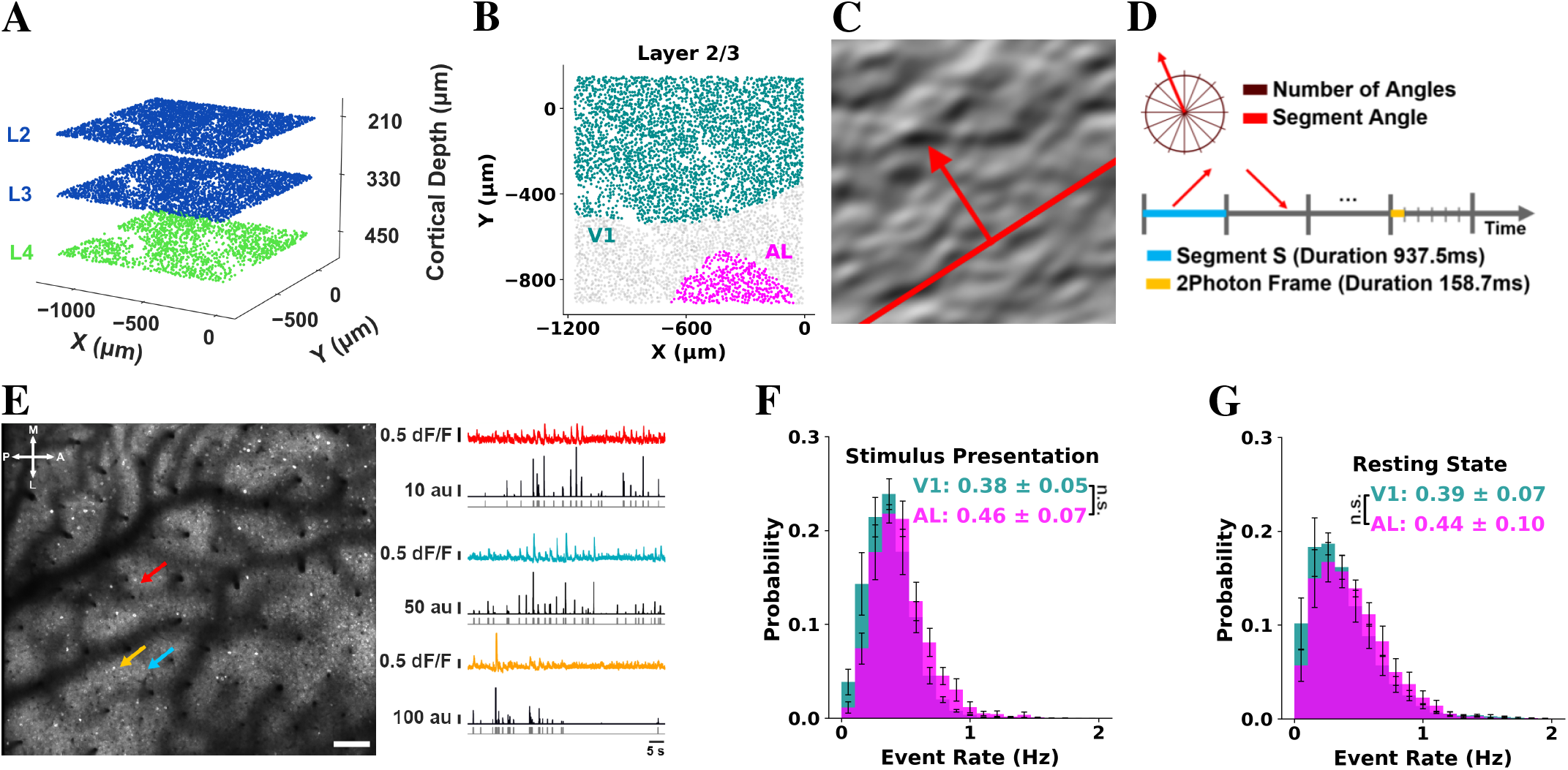
Imaging paradigm and visual stimulus. **A**. Illustration of supragranular-granular layer fields of view (FOVs) simultaneously acquired at ∼6Hz. L2/3: blue. L4: green. **B**. Retinotopic map of the FOV acquired in layer 2/3. V1: turquoise, AL: magenta, other areas: gray. Note that neurons within 15*µm* of the edge of the FOV or with an event rate below 0.01Hz during the visual presentation period have been excluded. In V1, the number of neurons was 917 ± 225 in layer 4 and 2960 ± 701 in layer 2/3. In AL, the number of neurons was 172 ± 103 in layer 4 and 485 ± 172 in layer 2/3. We report the mean and standard deviation across mice (n=5). **C**. Example frame of “Monet” video, consisting of waves with 16 distinct randomly shuffled directions of motion, presented to the mice (i.e., stimulus). The red arrow indicates the direction of motion of the stimulus. **D**. Example of a sequence of segments, each segment with a fixed stimulus direction. **E**. Example FOV acquired in layer 2/3 at depth 210*µm*. A: anterior, L: lateral, P: posterior, M: medial. Bar = 75*µm*. Color arrows indicate 3 example cell bodies whose traces are shown in color on the right. dF/F: fractional fluorescence change. au: the relative probability of firing in arbitrary units. Deconvolved firing probability traces shown in black below were obtained using the CaImAn algorithm [18]. The results are not sensitive to the threshold chosen. Gray traces in the bottom represent the thresholded, binarized, probability that specific imaging frames contain a calcium event (0: no event; 1: event). **F**. Pyramidal neuron event rate histogram of L2/3 neurons across animals during stimulus presentation. **G**. As in F, under resting state. Error bars represent the standard error of the mean across animals (n=5). Inset: Mean event rate ± standard deviation across animals (n=5), for each area. “n.s.”: non statistically significant. The highest p-value obtained from the permutation of means, the Welch’s t-test, and the ANOVA F-test is considered for the level-of-significance.

For each mouse, we obtained approximately *60-minute neuronal recordings*, during which mice were presented with stimuli videos of smoothened Gaussian noise with coherent orientation and motion, consisting of waves with **16 distinct** randomly shuffled directions of motion [20] (Figs. 1C & 1D, see Appendix), as well as a period of equal duration during which the mice were not presented with any stimulus (resting state). The 2-photon (2P) imaging recordings were preprocessed for motion correction and underwent automatic segmentation, deconvolution, and appropriate thresholding [18] to yield calcium “*eventograms*” that were analyzed (Fig. 1E, methods in Appendix). Neurons with event rates less than 0.01Hz or located less than 15*µm* from the field of view border are excluded.

The event rates of layer 2/3 neurons during the visual stimulation and resting-state periods are presented in Figs. 1F & 1G, respectively. Consistent with prior findings [7], the average event rates of neurons in V1 and AL show no statistically significant difference.

## III. Functional Connectivity

To identify functional connectivity patterns within the recorded layers, we applied the Spike Time Tiling Coefficient (STTC), a pairwise functional connectivity measure that exhibits several advantages over several other correlation measures (e.g., depends less on the firing rate than Pearson correlation) [3].

To quantify the temporal correlation between the firing events of a neuronal pair A and B, we employed their calcium eventograms and estimated the **STTC** weight as follows:

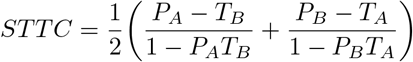

where *T*_*A*_ is the proportion of the recording duration that corresponds to firing events of neuron A, *P*_*A*_ the proportion of firing events of neuron A *synchronous* (i.e., within the same 158.7 ms frame) with firing events of neuron B, and likewise for *T*_*B*_ and *P*_*B*_. The STTC weight (correlation) takes values in [-1, 1]. To determine whether there is a *statistically significant functional connection between two neurons*, their STTC value was compared to a null distribution of STTC values calculated by circularly shifting their calcium eventogram time series by a random number, 500 times, yielding a z-score that determines the *level of significance*. We consider that two neurons are *functionally connected* when their z-score is *above 4*. Note that given the temporal kinetics of calcium imaging and the frame duration in our datasets (≈ 158.7 ms), the duration of a single frame is large compared to the neuron communication time (few ms).

We computed the STTC between all neuronal pairs belonging to the same layer (L2/3 and L4) within areas V1 and AL during visual stimulation (Figs. 2A & 2E). For both cortical layers, neuronal pairs in area AL exhibit stronger pairwise temporal correlations than those in V1. This difference is statistically significant. Additionally, AL has a higher number of statistically significant connections than V1, regardless of the threshold chosen for the z-score (Figs. 2B & 2F). The intra-layer degree of connectivity of a neuron is the number of neurons with which it has statistically significant functional connections, normalized by the neuronal population of their layer and area (see Appendix). In both layers, the distribution of the degree of connectivity in AL appears to be uniform, which is not the case in V1 (Figs. 2C & 2G). Functionally connected groups in AL are more cooperative than in V1, as they exhibit higher clustering coefficients (Figs. 2D & 2H, see Appendix).

**Fig. 2:**
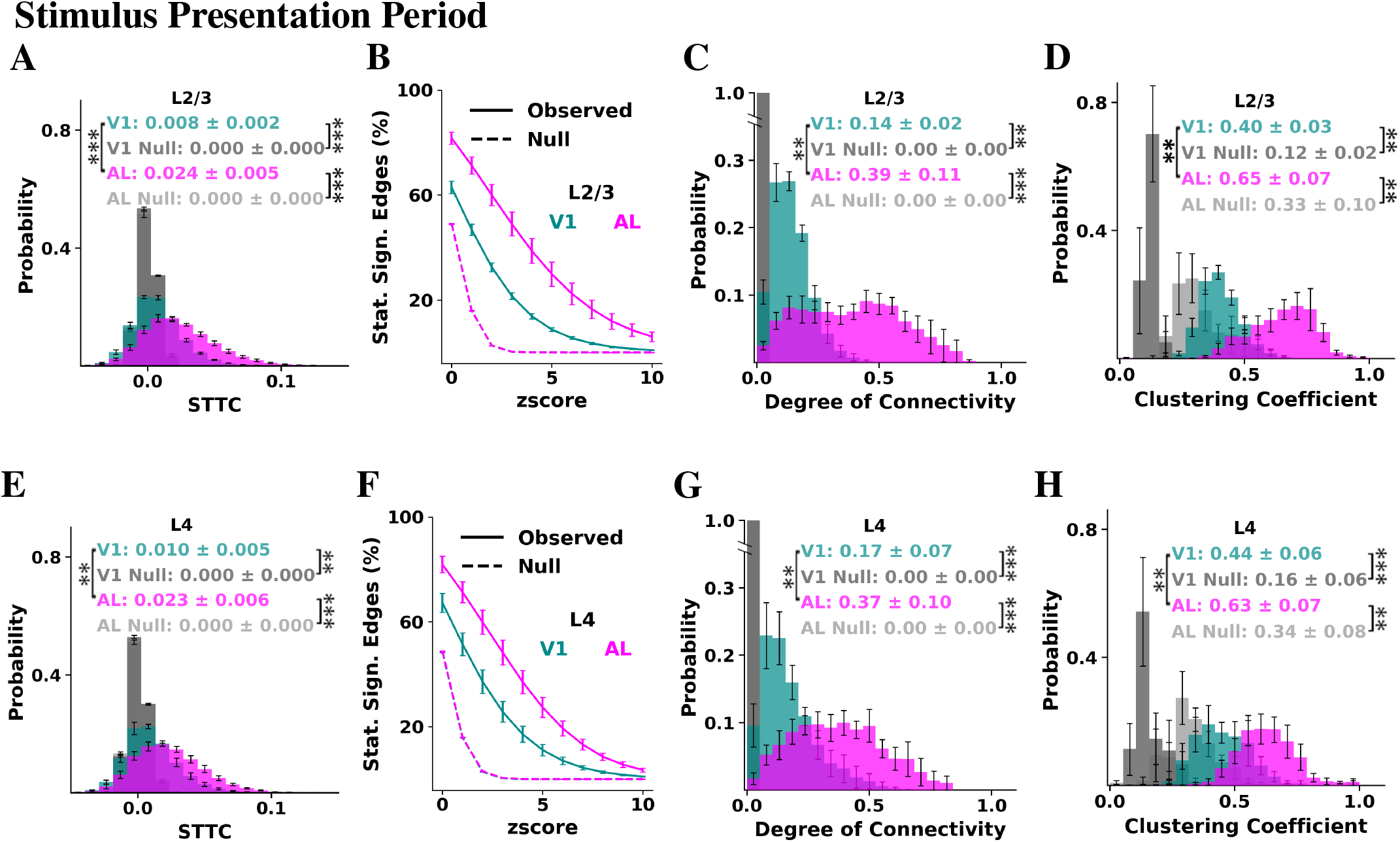
Functional connectivity of each cortical area during visual stimulation. **A**. Histogram of the STTC weights of all intra-L2/3 pairs for each cortical area, as computed during the stimulus presentation period. The control distributions were computed by randomly circularly shifting the eventograms. **B**. Percentage of intra-L2/3 connections identified as statistically significant as a function of the z-score threshold selected, as computed during the stimulus presentation period. The dashed lines indicate the controls, computed by randomly circularly shifting the eventograms. **C**. Histogram of the degree of connectivity during the stimulus presentation period considering intra-L2/3 connections (z-score>4). The control distributions were computed by randomly circularly shifting the eventograms. **D**. Distribution of clustering coefficients of L2/3 functionally connected groups during the visual stimulation period. The control distributions were computed on groups formed by randomly selecting neurons of the same layer and of equal size as the observed neuronal group of interest that does *not* overlap with it. **E-H**. Same as A-D, but considering only L4 neurons. Figure insets represent the mean ± standard deviation of the sample means across mice (n=5); Error bars represent the standard error of the mean across mice (n=5). P-values: “**”: < 0.01; “***”: < 0.001 The highest p-value obtained from the permutation of means, the Welch’s t-test, and the ANOVA F-test is considered for the level-of-significance.

We further characterized the functional connectivity during the resting state, i.e., in the absence of visual stimulation. The connectivity trends we observed under stimulus presentation largely persist (Fig. 3). However, a notable divergence emerges in L4, where *both* V1 and AL exhibit distributions of the degree of connectivity that are approximately uniform (Fig. 3G). When comparing each cortical area across stimulus presentation and resting state, we observed that within V1, the clustering coefficient exhibits a statistically significant increase during the resting state compared to the stimulus presentation period in both L2/3 (Figs. 2D & 3D, pvalue < 0.01) and L4 (Figs. 2H & 3H, pvalue < 0.01). Additionally, the STTC values in V1 show a statistically significant increase in L4 (Figs. 2E & 3E, pvalue < 0.05) during resting state compared to stimulus presentation. In contrast, we identified no statistically significant differences between the stimulus presentation and resting-state periods in the STTC values, degree of connectivity or clustering coefficient of area AL in either cortical layer. Thus, while certain functional connectivity metrics in V1 are dependent on the stimulation condition (visual presentation vs. resting state), in area AL they seem less affected by the condition than in V1. We report the highest p-value obtained from the permutation of means, the Welch’s t-test, and the ANOVA F-test.

**Fig. 3:**
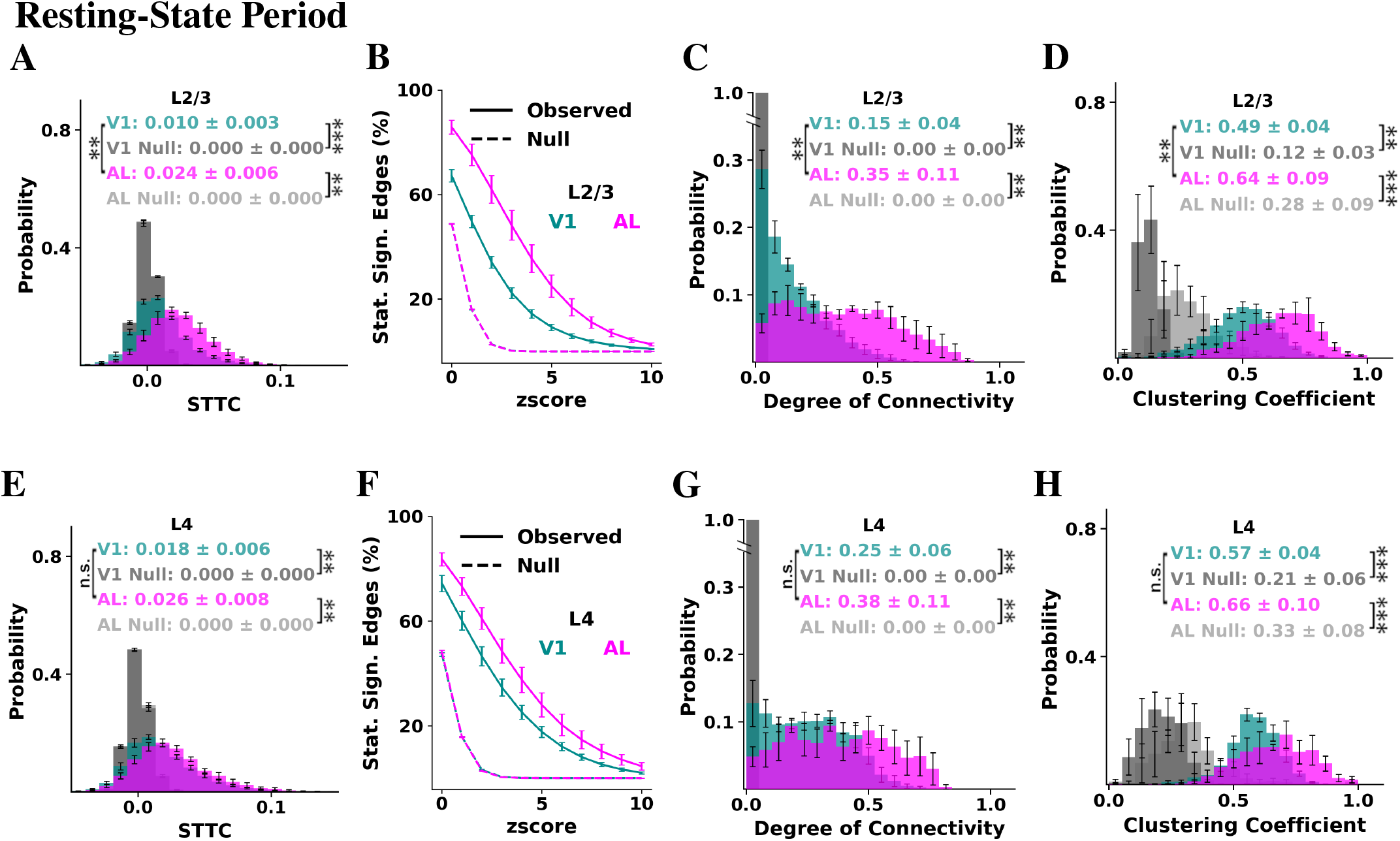
Functional connectivity of each cortical area during the resting state. **A**. Histogram of the STTC weights of all intra-L2/3 pairs for each cortical area, as computed during the resting-state period. The control distributions were computed by randomly circularly shifting the eventograms. **B**. Percentage of intra-L2/3 connections identified as statistically significant as a function of the z-score threshold selected, as computed during the resting-state period. The dashed lines indicate the controls, computed by randomly circularly shifting the eventograms. **C**. Histogram of the degree of connectivity during the resting-state period considering intra-L2/3 connections (z-score>4). The control distributions were computed by randomly circularly shifting the eventograms. **D**. Distribution of clustering coefficients of L2/3 functionally connected groups during the resting-state period. The control distributions were computed on groups formed by randomly selecting neurons of the same layer and of equal size as the observed neuronal group of interest that does *not* overlap with it. **E-H**. Same as A-D, but considering only L4 neurons. Figure insets represent the mean ± standard deviation of the sample means across mice (n=5); Error bars represent the standard error of the mean across mice (n=5). P-values: “**”: < 0.01; “***”: < 0.001, and “n.s.”: non statistically significant. The highest p-value obtained from the permutation of means, the Welch’s t-test, and the ANOVA F-test is considered for the level-of-significance.

## IV. Stability of Functional Connectivity Within and Across Stimulus Conditions

To evaluate the temporal stability of functional connections within each period, we divided the stimulus presentation and resting-state one-hour recordings into four consecutive 15-minute intervals. For each interval, we computed the statistically significant functional connections. We classified a connection as *persistent* if it remains statistically significant in each of the four individual intervals. This analysis revealed that, during both visual stimulation and resting state, L2/3 in area AL exhibits a statistically significantly higher proportion of persistent connections than V1 L2/3 (Fig. 4), suggesting higher temporal robustness of functional connections in AL compared to V1 during both conditions.

**Fig. 4:**
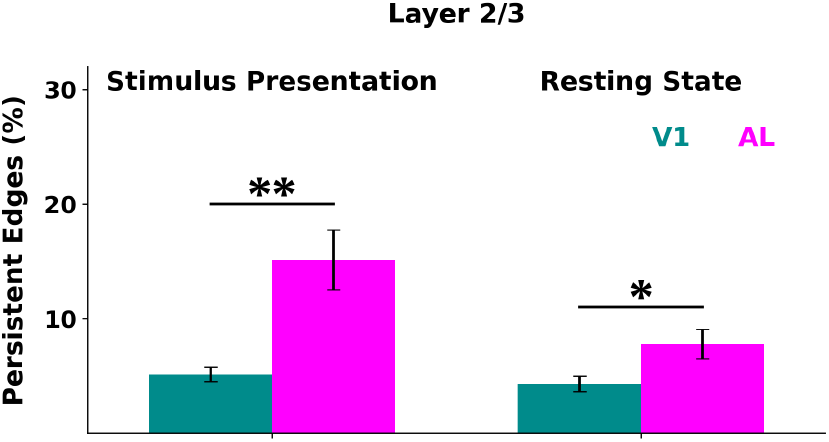
Statistically significant functional connections are more persistent in AL than in V1. Percentage of intra-L2/3 statistically significant functional connections identified as persistent during the stimulus presentation period (left) and the resting-state period (right) in V1 (turquoise) and AL (magenta). Each one-hour recording was divided into four consecutive 15-minute sub-periods. A functional connection is identified as persistent if it is statistically significant (z-score > 4) in each of the four individual periods. The percentage is calculated as the ratio of persistent connections to the total number of statistically significant connections detected over the entire period. All persistent connections are statistically significant when assessed across the full one hour recording. P-values: “*”: < 0.05 and “**”: < 0.01. The highest p-value obtained from the permutation of means, the Welch’s t-test, and the ANOVA F-test is considered for the level-of-significance.

To assess the stability of the functional connectivity between stimulus presentation and the resting state, we quantified the overlap of statistically significant edges (SSE) between the visual stimulation and resting-state conditions, normalized by the geometric mean of the SSE counts in each period. In both cortical layers, area AL exhibits a statistically significant higher overlap of functional connections present across both stimulus presentation and resting-state conditions compared to V1 (Fig. 5A). Additionally, we compared the degree of connectivity distributions between the stimulus presentation vs. resting-state periods. We discretized these distributions using a fixed bin size and contrasted them via the Jensen–Shannon Divergence (JSD) [21], which quantifies dissimilarity between probability distributions (see Appendix). For L4, the JSD between the visual stimulation and resting state distributions is higher in V1 than in AL (Figs. 5B & 5C), indicating a greater reorganization of V1’s connectivity between conditions compared to that of AL. This finding is robust to changes in bin size (Fig. 5D). Comparing the distributions of the degree of connectivity during the stimulus presentation and resting-state conditions in L2/3 yields no statistically significant differences between the two cortical areas.

**Fig. 5:**
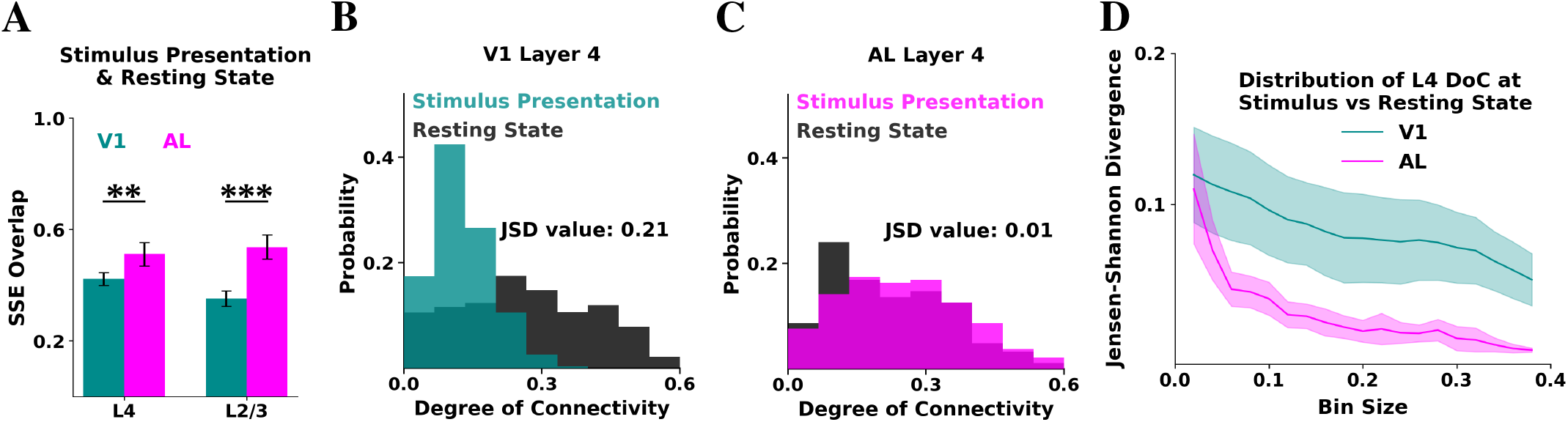
AL functional connectivity exhibits higher degrees of stability than V1 between the stimulus presentation and resting state periods. **A**. Overlap of statistically significant intra-L4 (left) and intra-L2/3 (right) edges between stimulus presentation and resting state for each cortical area. Overlap is quantified as the number of edges that are significant during both periods, normalized by the geometric mean of SSE counts in each period. **B**. Histogram for an example mouse of the intra-L4 degree of connectivity in V1 under stimulus presentation (turquoise) and resting state (black). The distributions were discretized using a bin of size 0.06. The Jensen-Shannon Divergence between the two distributions is reported. **C**. Same as B, but for AL of the same mouse. **D**. Jensen-Shannon Divergence between the distribution of the intra-L4 degree of connectivity during stimulus presentation vs resting state for each area. The x-axis represents the bin size used for discretizing the distributions. The lines and shaded regions correspond to the mean and SEM across mice, respectively. The difference is statistically significant for bin size = 0.3 (pvalue < 0.05). P-values: “**”: < 0.01 and “***”: < 0.001. The highest p-value obtained from the permutation of means, the Welch’s t-test, and the ANOVA F-test is considered for the level-of-significance.

## V. Dependence of Functional Connectivity on Pupil Size

We then examined how the behavior, using the pupil size as a proxy for arousal [16], [19], affects the activity and the functional connections of V1 and AL during the resting state. We computed the calcium event-triggered averages (ETAs) of V1 L2/3 neurons to the normalized pupil radius and found that V1 neurons tend to fire near the pupil size nadir (negative peak of the ETA) (Fig. 6A, turquoise line). Compared to V1 L2/3, AL L2/3 exhibits statistically significant weaker modulation by the pupil size (Fig. 6A, magenta line).

**Fig. 6:**
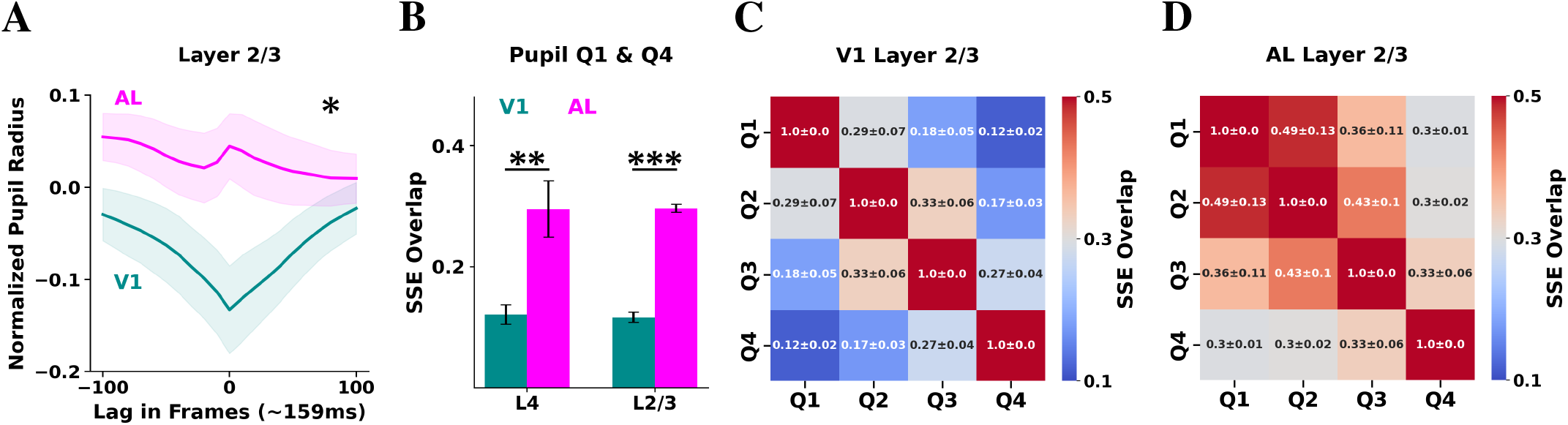
The activity and functional connections in AL are less dependent on pupil radius compared to V1 under resting state. **A**. ETA on pupil radius of L2/3 neurons for each cortical area, computed during the resting state. The lines and shaded regions indicate the mean and standard error of the mean across mice, respectively. Time is measured in imaging frames (158.7 ms) from the time of a calcium event fired by the L2/3 neuron (time zero). To evaluate the statistical significance of the difference between areas, we compare the ETA values at zero lag. **B**. Overlap of statistically significant intra-L4 (left) and intra-L2/3 (right) edges between periods of small (quartile 1) and large (quartile 4) pupil radius for each cortical area. Overlap is quantified as the number of edges significant during both periods, normalized by the geometric mean of SSE counts in each period. **C**. Heatmap illustrating the overlap of statistically significant intra-L2/3 edges between all pairs of pupil size quartile periods. Each cell in the heatmap shows the mean and standard deviation across mice of the overlap between the two periods. **D**. Same as C, but for AL. P-values: “*”: < 0.05, “**”: < 0.01 and “***”: < 0.001. The highest p-value obtained from the permutation of means, the Welch’s t-test, and the ANOVA F-test is considered for the level-of-significance.

To quantify the impact of arousal on network-level functional connectivity, we computed the overlap of the statistically significant connections between periods of small and large pupil radius. Specifically, we calculated the distribution of the pupil radius values and identified the imaging frames corre-sponding to the first (Q1) and fourth (Q4) quartiles. We then computed the statistically significant functional connections during each period. The AL network exhibits substantially higher SSE overlap between periods of small and large pupil radius than V1 (Fig. 6B), suggesting that its functional connectivity remains more stable under different arousal states. Extending this analysis to all pairwise comparisons of pupil radius quartiles, we observed consistently greater overlap in AL compared to V1 (Figs. 6C & 6D), indicating that AL maintains less arousal-dependent functional connections than V1.

## VI. Spatial Locality Bias in Functional Connectivity

The functional connectivity does exhibit spatial locality, e.g., significant positive pairwise correlations peaked within 300µm 19-34% [14]. So we wondered whether the observed differences in functional connectivity and stability between AL and V1 could be explained by the shorter average interneuronal distances in AL compared to V1. To address this, we first quantified the proportion of statistically significant connections as a function of distance. Even when restricting the analysis to edges of similar approximate length, AL tends to show denser functional connectivity than V1 under both stimulus presentation and resting-state conditions (Figs. 7A & 7B). We then computed the percentage of persistent edges as a function of inter-neuronal distance. In both V1 and AL, shortrange connections are more likely to be persistent compared to long-range ones (Fig. 7C). Notably, when restricting the analysis to short-range connections (< 200*µm*), AL shows a higher percentage of persistent edges than V1. We then com-puted the overlap of statistically significant edges across the stimulus presentation and resting-state periods as a function of inter-neuronal distance. In both V1 and AL, short-range connections show higher overlap than long-range ones (Fig. 7D). However, even when restricting the analysis to edges of similar length, AL exhibits higher overlap than V1. We extended this analysis to compare connectivity across epochs of small and large pupil radius and observed similar patterns to those we observed when comparing the stimulus presentation and resting-state conditions (Fig. 7E). These findings suggest that spatial locality bias does not fully account for the higher density, temporal robustness and cross-condition stability of the functional connectivity in AL compared to V1.

**Fig. 7:**
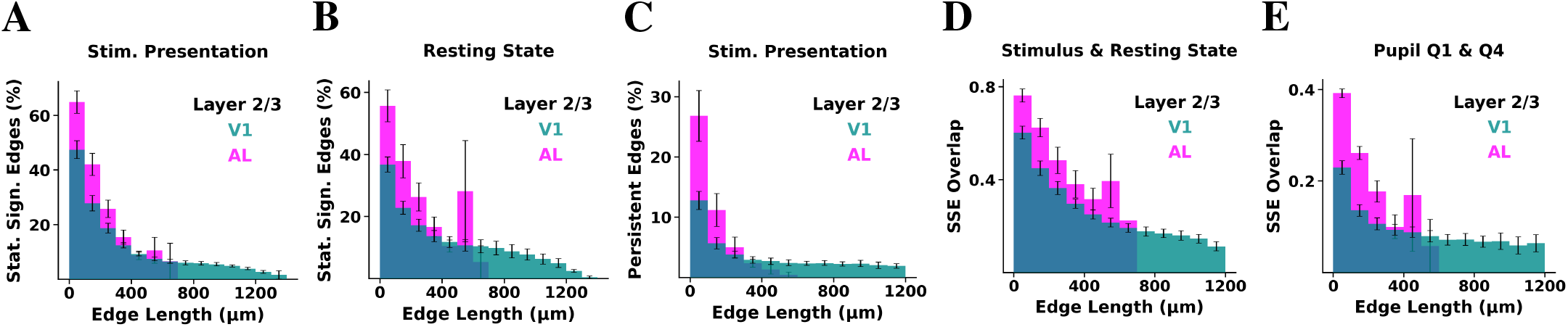
Spatial locality bias does not fully account for the higher temporal robustness and cross-state stability of the functional connections in AL compared to V1. **A**. Percentage of intra-L2/3 statistically significant edges (z-score > 4) as a function of distance between neuronal pairs, computed during the stimulus presentation period, plotted in bins of 100*µm*. The percentage is computed by taking as a denominator the number of all possible edges that could form at that distance between neurons of the same area (V1: turquoise, AL: magenta). **B**. Same as A, but computed during the resting-state period. **C**. Percentage of statistically significant edges that are identified as persistent as a function of distance between neuronal pairs, computed during the stimulus presentation period. **D**. Overlap of the statistically significant edges between the stimulus presentation and resting-state periods as a function of distance between neuronal pairs. To compute the overlap, we first select the edges whose lengths fall within the bin of interest. We then compute the number of edges within that bin that are statistically significant during both the stimulus presentation and resting-state periods, normalized by the geometric mean of the number of statistically significant edges in each period. **E**. Overlap of the statistically significant edges between periods of small (quartile 1) and large (quartile 4) pupil size, as a function of distance between neuronal pairs. Error bars represent the standard error of the mean across animals (n=5).

## VII. Discussion and Future Work

Our study reveals that the anterolateral (AL) visual area maintains a relatively stable functional connectivity across both stimulus-driven and resting-state conditions, compared to V1. The functional connectivity in AL consistently exhibits lower dependence on the presence of the visual stimulus and the arousal state, suggesting it is a more state-resilient cortical area than V1.

AL has a greater percentage of persistent connections over time, indicating that its functional connections remain robust across time than those in V1, both during stimulation and spontaneous activity, suggesting that AL’s network exhibits a “core” subnetwork that remains stable.

The quantification of the changes in functional connectivity between the stimulus presentation and resting-state periods, as assessed using the overlap of the SSE and the Jensen-Shannon Divergence metrics, revealed smaller differences in the structure of the functional connectivity between the stimu-lus presentation and resting-state conditions in AL than in V1. This suggests that the functional connectivity derived from spontaneous activity more faithfully reflects the functional network architecture elicited by visual stimulation in AL compared to V1.

We further demonstrated that pupil radius, a proxy of arousal [19], had a weaker modulatory effect on AL activity and functional connectivity compared to V1. L2/3 neurons in AL displayed nonsignificant modulation by the size of the pupil, and the overlap in statistically significant edges across pupil size quartiles remained consistently higher in AL than in V1.

It is part of our ongoing research to examine the temporal robustness and stability of other layers, as well as other higher-order visual circuits, namely the lateromedial (LM) and rostro-lateral (RL) areas, and assess whether stability systematically increases along the visual cortical hierarchy. Furthermore, the impact of different types of stimuli on the functional connectivity of visual areas requires further exploration.

## Acknowledgments

We extend our sincere appreciation to Taliah Muhammad of the Tolias Laboratory (formerly at Baylor College of Medicine, now at Stanford University) for her expertise in performing the two-photon mesoscope experiments. We are also grateful to other members of the Tolias Laboratory, particularly Paul Fahey, for their valuable feedback on the data. We extend our special thanks to Andreas Tolias for his generous collaboration and support throughout this project. We would also like to thank Ganna Palagina for preprocessing the data and developing the data pipeline, and to Ioannis Smyrnakis and Georgios Keliris for insightful discussions on various aspects of the analysis. We are also grateful to the data analysts at FORTH, Nikolaos Tzanakis, Manos Koniotakis, and Vassilis Sideridis, who contributed to the preliminary analysis.

This work has received funding from the European Union’s Horizon 2020 research and innovation program under the Marie Skłodowska-Curie grant agreement No 101007926 as well as from the Hellenic Foundation Research Institute (HFRI) with the neuron-AD project number 04058 and neuronXnet project number 2285 (PI: Maria Papadopouli). It has been partially supported by project MIS 5154714 of the National Recovery and Resilience Plan Greece 2.0 funded by the European Union under the NextGenerationEU Program. Finally, this research was also supported by R01 NS113890, and R21 NS127299 (PI: Stelios Smirnakis).

APPENDIX

### Mouse Lines and Surgery

Five adult mice (10-12 weeks of age), expressing GCaMP6s in excitatory neurons via SLC17a7-Cre and Ai162 transgenic lines, were anesthetized and a 5mm craniotomy was placed over visual cortex as described [22]. Each mouse recovered for ∼2 weeks prior to the first experimental imaging session.

#### Experimental Data Collection

The animals underwent mesoscopic two-photon imaging covering most of dorsal area V1 and nearby extrastriate cortex, while being head-fixed on a treadmill in quiet wakefulness. Images were acquired at 6.30072 Hz over a ∼1.2×1.2 mm^2^ field of view sampling simultaneously across 4 planes corresponding to V1 layers 2 (80-210 *µm*), 3 (285-330 *µm*), 4 (400-450 *µm*) and 5 (500 *µm*). Images were preprocessed in standard fashion for motion correction and underwent automatic segmentation and deconvolution using the CNMF CaImAn algorithm [18]. The deconvolved signal was thresholded to yield calcium “eventograms” that were used for analysis. The threshold yielding calcium event rates closest to those reported in the literature [23] was selected.

#### Monitor Positioning and Retinotopy

Visual stimuli were presented to the left eye with a 31.1×55.3cm^2^ (h×w) monitor (resolution of 1440×2560 pixels) positioned 15cm away from the mouse eye. Pixelwise responses across a 2400×2400 *µ*m^2^ to 3000×3000 *µ*m^2^ region of interest (0.2 px/*µ*m) at 200-220*µ*m depth from the cortical surface to drifting bar stimuli were used to generate a sign map for delineating visual areas [24].

#### Directional Visual Stimulus

A stimulus using smoothened Gaussian noise with coherent orientation and motion was presented to the mice. An independently identically distributed (i.i.d.) Gaussian noise movie was passed through a temporal low-pass Hamming filter (4Hz) and a 2-d Gaussian filter (*σ* = 4.4^*°*^ at the nearest point on the monitor to the mouse). Each scan contained 72 blocks, with each 15-second block comprising of 16 equally distributed and randomly ordered unique directions of motion between 0-360 degrees with a velocity of 42 degrees/s at the nearest point on the monitor. An orientation bias perpendicular to the direction of movement was imposed by applying a bandpass Hanning filter G(*ω*; c) where *ω* is the difference between the image 2d Fourier transform polar coordinates *ϕ* and trial direction *θ*, and

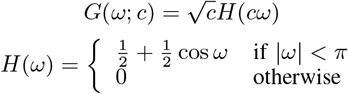

Here, c = 2.5 is an orientation selectivity coefficient. The resulting kernel is 72^*°*^ full width at half maximum.

#### Normalized Degree of Connectivity

For each neuron, we estimate the fraction of neurons with statistically significant functional connections (z-score > 4) to that neuron, per layer case, which corresponds to the normalized degree of connectivity in that layer. For example, a L4 neuron has an intra-layer degree of connectivity of 0.1 if that neuron is functionally connected with 10% of the L4 neuronal population.

#### Clustering Coefficient

For each neuron, we estimate the fraction of neuronal pairs of its neighbors that form statistically significant functional connections with each other. For example, an L2/3-neuron with an intra-L2/3 clustering coefficient of 10% indicates that exactly 10% of all possible pairwise functional connections between its L2/3 neighbors are statistically significant.

#### Jensen-Shannon Divergence

To quantify differences between discrete probability distributions of intra-layer degree of connectivity under different conditions, we computed the Jensen–Shannon Divergence (JSD) [21]. Given two distributions *P* and *Q*, JSD is defined as:

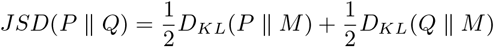

where 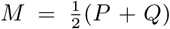 is the average distribution, and *D*_*KL*_ denotes the Kullback–Leibler divergence:

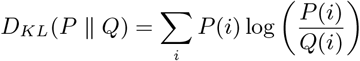

JSD is symmetric, bounded between 0 and 1 (when using base-2 logarithm), and well-suited for comparing empirical probability distributions derived from discretized connectivity metrics. The distributions were normalized histograms discretized with fixed bin sizes.

